# Cooperation genes are more pleiotropic than private genes in the bacterium *Pseudomonas aeruginosa*

**DOI:** 10.1101/2022.06.25.495533

**Authors:** Trey J. Scott

**Author notes:** Corresponding author: Trey J. Scott **Email:**. **Author Contributions:** TS designed the study, analyzed the data, and wrote the manuscript.

## Abstract

Pleiotropy may affect the evolution of cooperation by limiting cheater mutants if such mutants also lose other important traits. Because pleiotropy limits cheaters, selection may favor cooperation genes that are more pleiotropic. However, the same should not be true for private genes with functions unrelated to cooperation. Pleiotropy in cooperative genes has mostly been studied with single genes and has not been measured on a wide scale or compared to a suitable set of control genes with private functions. I remedy this gap by comparing genomic measures of pleiotropy in previously identified cooperative and private gene sets in *Pseudomonas aeruginosa.* I found that cooperative genes in *P. aeruginosa* tended to be more pleiotropic than private genes according to the number of protein-protein interactions, the number of gene ontology terms, and gene expression specificity. These results show that pleiotropy may be a general way to limit cheating and that cooperation may shape pleiotropy in the genome.

## Introduction

Many genes are pleiotropic, where a pleiotropic gene is defined as one that affects multiple traits. Pleiotropy constrains evolution because mutations with beneficial effects on one trait can have deleterious effects on other traits. Several measures of pleiotropy have been used and shown to be associated with evolutionary constraint. Three examples are the number of protein interactions (1), the number of functional annotations (2), and how widely genes are expressed across tissues (3).

Pleiotropy is thought to limit cheaters and stabilize cooperation (4–7; see 8 for an alternate way pleiotropy can affect cooperation and 9 for a dissenting view). Explaining how cheater evolution can be limited is a central question in the study of cooperation (10). Cheaters benefit from cooperation without paying the costs and are expected to outcompete cooperators. This cheater advantage can lead to the breakdown of cooperation unless cheaters are constrained (10).

Pleiotropy can limit cheaters when mutations that cause cheater phenotypes come with harmful effects on other traits. One example of this comes from the social amoeba *Dictyostelium discoideum. D. discoideum* has a cooperative stage where 20% of cells sacrifice themselves to become stalk. The remaining cells become spores that are held up for dispersal by the stalk (10). This act of cooperation can be exploited by cheaters that contribute less to the stalk and increase their abundance in spores (10). Amoebae with *dimA* mutations are potential cheaters because they ignore the signal to become stalk (4). This should increase *dimA* mutant representation as spores. Instead, *dimA* mutants are excluded from spores when mixed with wildtype cells as a pleiotropic effect (4). This trade-off between becoming a stalk cell and entering spores could limit the cheating ability of *dimA* mutants.

Because pleiotropy limits cheaters, it should be higher in cooperative genes relative to private (non-cooperative) genes (5). It is unknown whether cooperative genes are generally more pleiotropic than private genes. Prior studies of pleiotropy in cooperative genes mostly focused on individual genes (4) or on the observation that genes are co-regulated (7). Such studies do not explicitly quantify pleiotropy or compare between cooperative and private genes. Private genes are an important control because they should reflect the background pleiotropy that is not affected by social selection.

Here, I take a genomic approach to compare cooperative and private genes as categorized by Belcher et al. (11) in four *P. aeruginosa* gene sets. These gene sets have been constructed so that cooperative and private genes in a set are expressed under similar conditions. Because of this, cooperative and private genes should be similarly exposed to selection, all else being equal within a set (11). These sets of cooperative and private genes are therefore ideal for testing whether cooperative genes are more pleiotropic.

## Results

To test whether pleiotropy is higher in cooperation genes compared to private genes, I first used 315 quorum sensing genes (12) that were previously categorized as cooperative (N = 41) or private (N = 274) based on gene annotations and experimental data (11). As measures of pleiotropy, I used the number of protein interactions contained in the STRING database (13), the number of biological process Gene Ontology (GO) terms (14), and gene expression pleiotropy for each gene with available data (Figure 1). These pleiotropy measures were not highly correlated and thus represent independent measures of pleiotropy (see appendix).

**Fig. 1.**
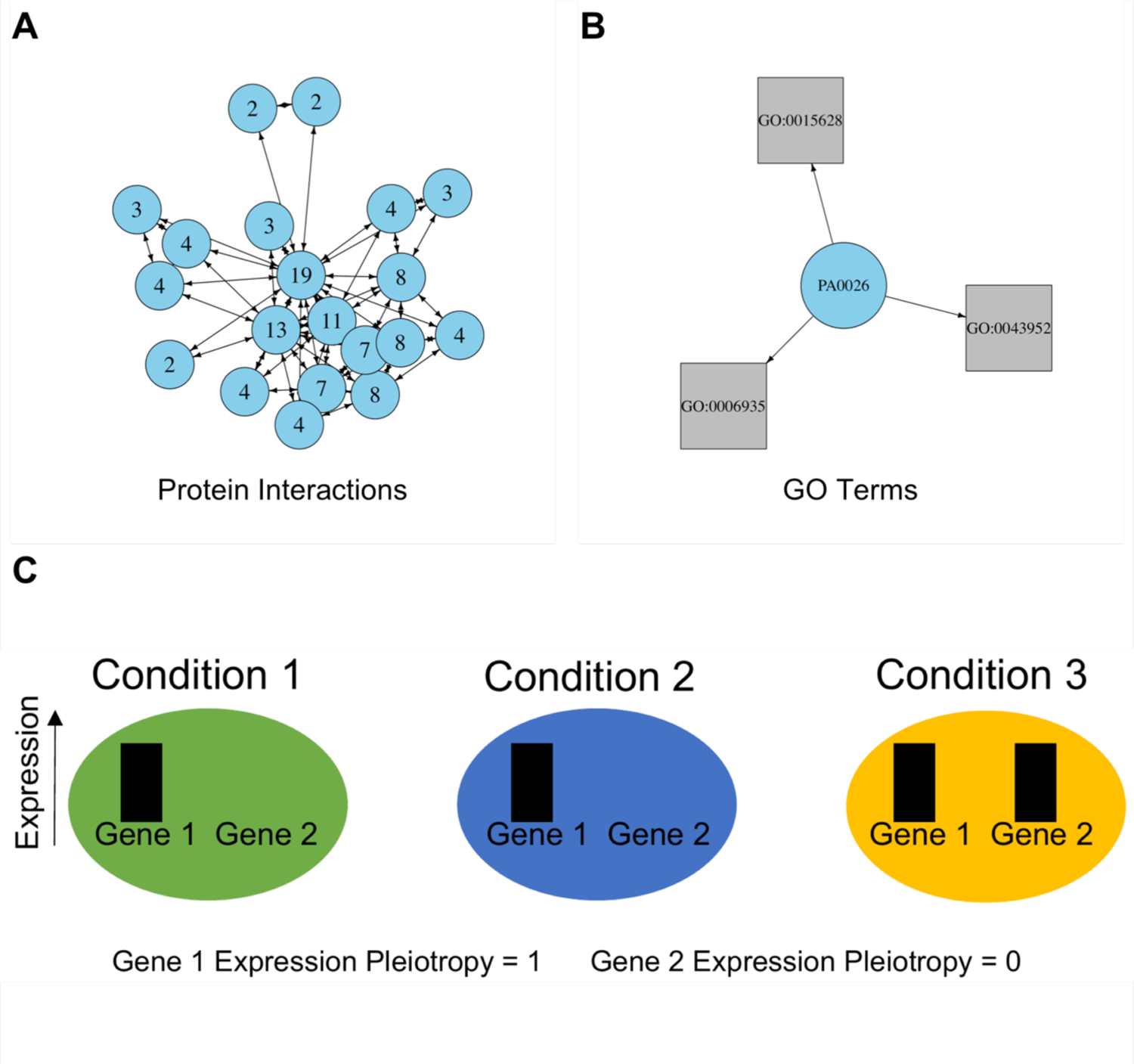
Examples of pleiotropy measures used in this study. (A) Number of protein interactions (STRING database). Arrows show predicted interactions that are summed to measure pleiotropy (the numbers on the nodes). (B) Number of biological process Gene Ontology (GO) terms from the *P. aeruginosa* genome database. Arrows show annotations for an example gene with three GO terms. (C) Gene expression pleiotropy is calculated from gene expression data across multiple conditions. Gene 1 is a gene that is expressed in every condition and is maximally pleiotropic. Gene 2 is a gene that is specialized for a single condition and is minimally pleiotropic.

I found that cooperative genes in the quorum sensing pathway were more pleiotropic across all three measures of pleiotropy (Fig. 2A). Cooperative genes had about 65 more STRING interactions on average than private genes (generalized linear model (GLM); p = 0.0162). Cooperative genes had 2 GO terms while private genes tended to have only 1 (GLM; p = 0.0090). Gene expression pleiotropy was around 15% higher in cooperative genes than private genes (beta regression; p = 0.0011).

**Fig. 2.**
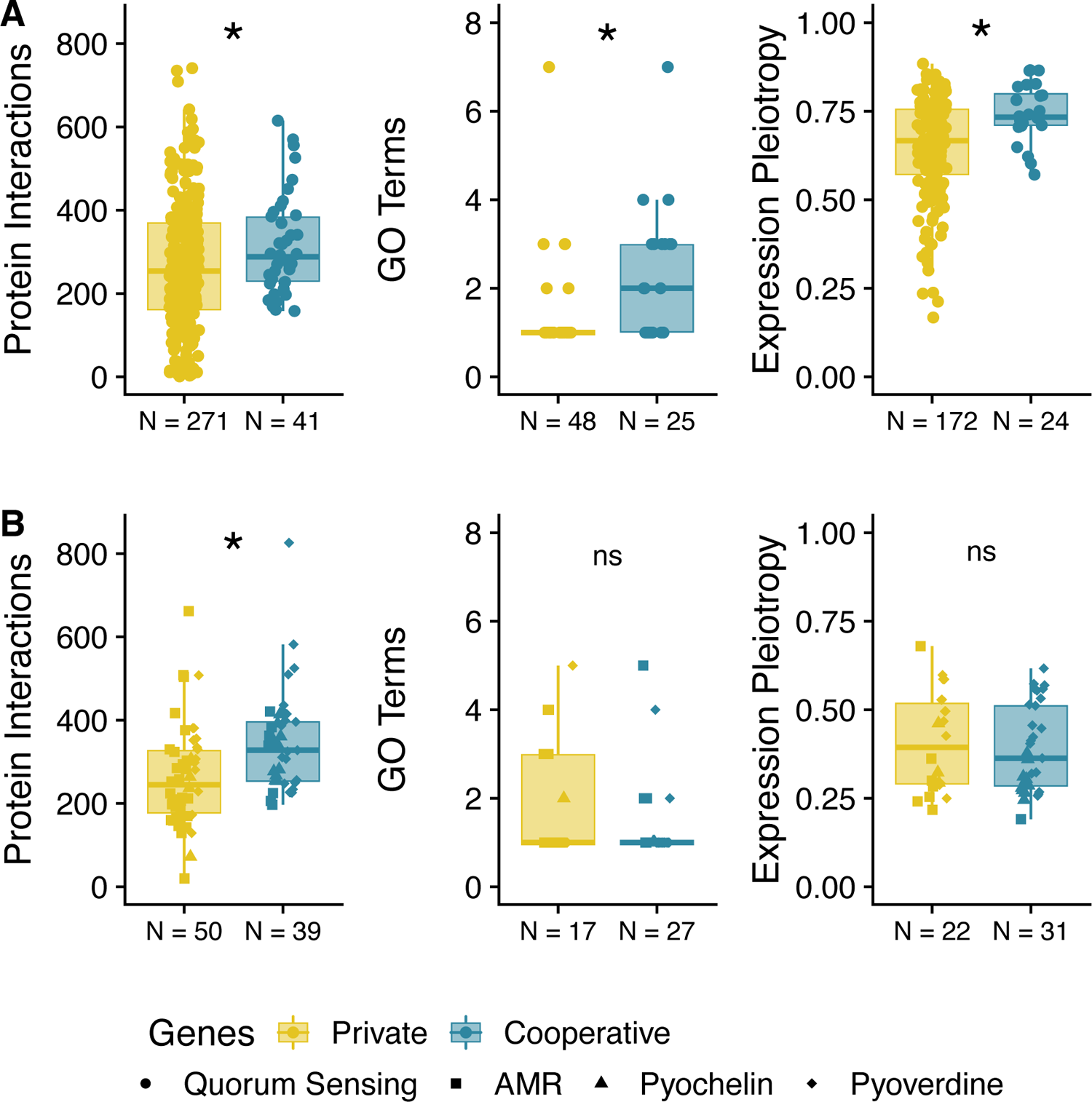
Cooperative genes tend to be more pleiotropic than private genes in the (A) quorum sensing and (B) other gene sets in *P. aeruginosa* (AMR = antibiotic resistance). Panels show the number of protein-protein interactions in the STRING database (left), the number of gene ontology (GO) terms (middle), and the gene expression pleiotropy (right). Number of genes with measures of pleiotropy are shown on the x-axis. Colors are the same as in (10) for easy comparison. *’s indicate p ≤ 0.05.

To test whether these results apply beyond the quorum sensing pathway, I tested additional sets of cooperative and private genes in *P. aeruginosa* that were included in Belcher et al. (11). These gene sets consisted of antibiotic resistance genes (AMR) and pyochelin and pyoverdine genes that are involved in binding iron. To increase statistical power, I analyzed all three gene sets together (Fig. 2B; see appendix). Cooperative genes tended to have more STRING protein interactions than private genes across these three gene sets (GLM; p = 0.0078). However, cooperative and private genes were not different in terms of GO terms (GLM; p = 0.371) and expression pleiotropy (beta regression; p = 0.102).

## Discussion

Cooperation can break down because of the evolution of cheaters that benefit from cooperation without helping (10). The advantage of a cheater mutation can be limited if it also has pleiotropic effects on other important traits (4–7). This may result in high pleiotropy in cooperative genes to limit cheaters. Prior studies have focused on single genes (4) and coregulation (7) instead of measuring pleiotropy across many genes.

Using three independent measures of pleiotropy in *P. aeruginosa*, I found that pleiotropy tended to be higher in cooperative genes than in private genes (Fig. 2). The majority of comparisons in this study supports the hypothesis that pleiotropy limits cheaters in populations of cooperators. In two comparisons, I found no difference between cooperative and private genes (Fig. 2B). However, these comparisons had the smallest number of genes with measures of pleiotropy and may have lacked power to detect a difference.

Another implication of increased pleiotropy in cooperation genes (Fig. 2) is a strengthening of the conclusions of Belcher et al. (11), who found evidence of kin selection because of relaxed selection in cooperation genes relative to private genes. Pleiotropy can also affect patterns of selection, but it should be in the opposite direction of kin selection. Pleiotropy is associated with signals of conservation (2) and stabilizing selection (15), which should decrease genetic diversity and divergence. This means that the signal of kin selection in (11) may be an underestimate since increased pleiotropy in cooperation genes should dampen the signal of relaxed selection

Theoretical studies have disagreed about the direction of causality between cooperation and pleiotropy and whether pleiotropy is able to stabilize cooperation if pleiotropy itself can evolve (5, 6, 8, 9). While the data in this study cannot resolve these theoretical disagreements, they show that pleiotropy and cooperation are linked in *P. aeruginosa*. Cooperation may thus shape patterns of pleiotropy in the genomes of other cooperative organisms through its link with high pleiotropy.

## Materials and Methods

I used cooperation and private genes identified in Belcher et al (11) and generated pleiotropy measures (see Figure 1, the appendix, and Datasets S1-S3). To test for differences between cooperative and private genes, I used generalized linear models (GLMs) and beta regression in R version 4.1.2. More detailed methods can be found in the appendix. Data and R code for analysis are available at www.gitlab.com/treyjscott/kin_selection_pleiotropy.

## Acknowledgments

I thank Laura Walker, Tyler Larsen, Joan Strassmann, and David Queller for comments on the manuscript. This work was supported by NSF awards DEB 1753743 and IOS-1656756.

## Notes

**Competing Interest Statement:** I have no competing interests to disclose.

https://gitlab.com/treyjscott/kin_selection_pleiotropy

### Competing Interest Statement

The authors have declared no competing interest.

### Summary of Updates

Re-uploaded as a pdf to fix bad rendering of Fig. 1.

