## appendix for "Cooperation genes are more pleiotropic than private genes in the bacterium *Pseudomonas aeruginosa*"

Trey J. Scott
Department of Biology, Washington University in St. Louis

Corresponding author: Trey J. Scott
Orcid ID: 0000-0001-6609-9638

**This PDF file includes:**

Appendix
SI References

**Other supplementary materials for this manuscript include the following:**

Datasets S1 to S3

**Appendix**

*Gene Sets*

I gathered 315 quorum sensing genes from Schuster et al. (1) and categorized 41 of these genes as cooperative from Belcher et al. (2). The remaining genes were classified as private genes that did not have a social function (Dataset S1). Additional pyochelin, pyoverdine, and AMR cooperative and private genes were also gathered from Belcher et al. (2) and are provided in Dataset S2.

*Pleiotropy*

I investigated three measures of pleiotropy (Dataset S3). First, I used predicted protein-protein interactions for *P. aeruginosa* PAO1 downloaded from the STRING database version 11.5 on May 6$^{th}$, 2022 (3). STRING contains predicted protein interactions collected from high-throughput experiments, text mining, and other resources. These interactions may not necessarily involve physical interactions between proteins but should convey functional relationships. STRING entries have confidence scores that provide a measure of quality for interaction predictions. I incorporated this measure in statistical models by weighting according to the average score for a protein's combined interactions.

My second measure of pleiotropy was the number of non-redundant biological process gene ontology annotations for *P. aeruginosa* PAO1. These annotations convey information about the functions that a gene has or is predicted to have (4). These data were downloaded on April 4$^{th}$ 2022 from the *P. aeruginosa* genome database version 20.2 (5), which updates annotations based on new results published on *P. aeruginosa*.

My final measure was gene expression pleiotropy, a measure of how widely genes are expressed across conditions. I calculated gene expression pleiotropy as 1 - τ, where τ is a common measure of gene expression specificity (6). τ ranges from 0, when a gene is expressed in all conditions tested, to 1, where the gene is expressed in only 1 condition and is calculated for each gene as

$$\tau = \frac{\sum_i \left( \left( 1 - ln(x_i) \right) / ln(x_{max}) \right)}{N-1},$$

where N is the number of conditions, $x_i$ is the expression level in conditions *i*, and $x_{max}$ is the maximum expression across all conditions (7). τ is usually calculated across different kinds of tissues. Since *P. aeruginosa* does not have conventional tissues, I instead used gene expression data from four timepoints, two during planktonic growth at 4 and 12 hours and two during biofilm growth at 24 and 48 hours (8). To ensure that log expression was positive, I manually changed the minimum expression to 1.

*Statistics*

To determine whether the three pleiotropy measures were independent, I checked for correlations using Spearman's ρ. Correlations were weak ranging from 0.019 between GO terms and expression pleiotropy to -0.238 between protein interactions and expression pleiotropy. The correlation between GO terms and protein interactions was 0.039. These pleiotropy measures were thus relatively independent.

To test whether cooperative genes were more pleiotropic than private genes for the quorum sensing pathway, I used generalized linear models (GLMs). For protein interactions and GO terms, I fit models with Poisson errors. If I detected overdispersion, I fit quasi-Poisson and negative binomial models for the final analysis. I conservatively reported the highest p-value between quasi-Poisson and negative binomial models if more than one model was fit. To include STRING confidence scores (see above), I weighted protein interaction models by the average score of its interactions. For gene expression pleiotropy, I used beta regression (9) to account for this measure being bounded from 0 to 1. To calculate means and p-values from statistical models, I used the *emmeans* package (10). I performed statistical tests in R (11) (version 4.1.2).

To test for differences between cooperative and private genes for the additional gene sets, I again used GLMs as above. I included the pathway (pyoverdine, pyochelin, or AMR) as an interaction term with sociality (private or cooperative) in models, but only compared means between cooperative and private genes (this averages over the effect of pathway).
